# Sex differences in vaccine-induced neuraminidase cross-recognition impact H5N1 dissemination to the lower respiratory tract in mice

**DOI:** 10.64898/2026.05.26.728011

**Authors:** Sabal Chaulagain, Anne P. Werner, Maclaine A. Parish, Sattya Narayan Talukdar, Anna Yin, Brittany A. Seibert, Tianle Zhang, Jennifer A. Liu, Cosette Schneider, Lynda Coughlan, Andrew Pekosz, Sabra L. Klein

## Abstract

H5N1 vaccines have been poorly immunogenic in humans, creating a challenge for vaccine development. Seasonal influenza vaccines offer some cross-protection against H5N1, but there has been no consideration of whether protection differs between the sexes. We investigated sex differences in antibody responses following receipt of either beta-propiolactone inactivated whole virus H1N1 or H5N1 (LAIV backbone) vaccines in C57BL/6 mice. Using systems serology assays, vaccination induced strong homologous and heterologous antibody responses, with females generating greater IgG titers than males against whole virus H1N1 and H5N1, which was primarily mediated by greater IgG responses to neuraminidase (NA) than hemagglutinin (HA) protein. Cross-reactive H5N1 IgG titers were greater among H1N1-vaccinated females, and primarily mediated by greater N1-specific IgG titers. IgG2b and IgG2c were the primary antibody isotypes generated in response to these vaccines, with females having greater IgG2b titers and enhanced binding to FcγRIV for avian and human NA than males following either homologous or heterologous vaccination. Antibody-dependent complement deposition was measured as an FcR-mediated non-neutralizing response against HA and NA and was more robust among H1N1 and H5N1 vaccinated females than their male counterparts in response to homologous HA only. Vaccinated females tended to have greater neutralizing antibody titers than males against the homologous vaccine strain, with limited cross-neutralizing antibodies detected in either sexes. Neuraminidase inhibition titers were greater in vaccinated females than males against the heterologous virus following H1N1 vaccination and against both the vaccine and heterologous viruses following H5N1 vaccination. When H1N1 and H5N1 vaccinated mice were challenged with a lethal dose of A/Texas/37/2024 H5N1, all H5N1 vaccinated mice were protected, regardless of sex. Among H1N1 vaccinated mice, while both sexes were protected against disease, H1N1 vaccinated females restricted virus to the upper respiratory tract and had lower pulmonary virus titers than males at 3 days post challenge. These findings highlight that sex differences in vaccine-induced NA-specific antibody responses are associated with differential respiratory dissemination of H5N1 and that sex should be considered in studies of vaccine-induced cross-reactive influenza immunity.

**Highlights:**

- Females mount stronger IgG responses than males to both H1N1 and H5N1 vaccines
- Sex differences in vaccine responses are driven by immunity to neuraminidase (NA)
- Females mount stronger anti-NA IgG2b responses with greater binding to FcγRIV than males
- NA inhibition titers are greater in females, supporting broader cross-immunity
- H1N1-vaccinated females contain H5N1 virus replication better than males after lethal challenge

## Introduction

Influenza viruses are classified by their surface glycoproteins, hemagglutinin (HA) and neuraminidase (NA), both of which are targets of protective antibody, although current inactivated vaccines are standardized on HA content alone (1–3). The A/California/07/2009 virus (H1N1), emerged by reassortment in 2009 and remains a component of seasonal vaccines (4). Highly pathogenic avian influenza A (H5N1) viruses of clade 2.3.4.4b have spread globally in birds since 2021 and were detected in US dairy cattle in 2024, causing a multistate outbreak with sporadic human cases, mostly following occupational exposure (5, 6). Population immunity to H5N1 HA is minimal, but 2009 H1N1 and clade 2.3.4.4b H5N1 share the N1 subtype of NA, raising the question of whether N1 immunity contributes to cross-recognition and possibility protection (7, 8).

Adult females produce greater neutralizing antibody responses than age-matched males following vaccination against either seasonal or pandemic influenza strains in humans (9, 10) and mice (11, 12). A meta-analysis of 33,092 healthy adults from 19 randomized controlled trials shows that the immunogenicity of influenza vaccines is greater in females compared to males (13). Even the breadth of immunity generated following receipt of an inactivated 2009 H1N1 vaccine is greater in females than males, at least among mice (12). Using a panel of 2009 H1N1 viruses that contain mutations in the immunodominant hemagglutinin (HA) protein, we previously demonstrated that following inactivated 2009 H1N1vaccination, adult females produce greater virus-specific, class-switched total IgG and IgG2c antibodies against the vaccine and all mutant viruses, and antibodies from females recognize more unique, linear HA epitopes, including from H5, than antibodies from males (12). Females also generate greater numbers of germinal center (GC) B cells that have greater somatic hypermutation (SHM) frequencies than vaccinated males (12). Deletion of activation-induced cytidine deaminase (*Aicda)* eliminates vaccine-induced female-biased immunity and protection against mutant viruses highlighting that SHM is necessary for the greater cross-protection in females (12). With H1N1 vaccinated females having greater breadth of immunity than vaccinated males, we hypothesize that cross-reactive immunity, including to NA, and protection against H5N1 may be similarly influenced by sex.

To date, studies in humans and male ferrets report limited cross-protection against H5N1 following intramuscular vaccination with seasonal inactivated influenza vaccines (IIV), with no detectable cross-neutralizing antibodies produced against H5N1 viruses (14). Intranasal vaccination with seasonal IIV combined with a toll-like receptor (TLR) 3 agonist induces cross-reactive binding antibodies and results in more rapid control of H5N1 virus replication following live virus challenge in female mice (15). It is well established that the quantity and quality of NA in conventional IIVs is not quantified (16). A meta-analysis of ferret studies further demonstrates that seasonal IIVs that do not contain neuraminidase (NA) subtype 1 (N1) provide limited cross-protection against H5N1, suggesting that N1 is desirable for cross-immunity against H5N1 (17). In humans, infection with H1N1, but not receipt of seasonal IIV, induces cross-reactive IgG, neutralizing antibody (nAb), and NA-inhibition (NAI) antibody titers against clade 2.3.4.4b H5N1, further highlighting that incorporation of NA is necessary for cross-recognition of H5N1 (7). At a population level, NAI titers against the N1 from clade 2.3.4.4b H5N1, but not HA-inhibition titers, are highly detectable (18). In female mice, NA-targeting monoclonal antibody (FNI9) protects against disease following challenge with clade 2.3.4.4b H5N1 (19). To date, however, whether the sex of the animal, patient, or vaccinee impacts the degree of cross-protection against H5N1 afforded by IIVs that contain N1 has not been reported and forms the basis of the current study.

## Results

### Females have greater homologous and heterologous virus-specific IgG titers than males

To determine if there were sex differences in the antibody isotypes generated in response to either beta-propiolactone (BPL) inactivated whole virus H1N1 or H5N1 vaccination, plasma IgM, IgA, and IgG were measured. At 35 days post vaccination (dpv), both IgM and IgA were produced at low titers against homologous HA and NA in H1N1 and H5N1 vaccinated mice, with no sex differences observed (**Fig. S1A-H**). Neither H1N1 nor H5N1 vaccinated mice produced detectable heterologous titers of IgM beyond very low titers observed in a small subset of animals and no detectable heterologous IgA responses against HA or NA were observed (**Fig. S1A-H**).

Consistent with previous studies (11, 12, 20, 21), after H1N1 vaccination, adult females had greater anti-H1N1 IgG titers against whole virus than males, as well as greater cross-reactive anti-H5N1 IgG titers against whole virus (**Fig. 1A**). After receipt of H5N1 vaccine, adult females developed greater anti-H5N1 IgG and cross-reactive anti-H1N1 IgG titers against whole viruses than males (**Fig. 1B**). Because the ELISA antigens were IIVs, IgG recognized multiple viral proteins, including conserved internal antigens (e.g., nucleoprotein and matrix protein 1), and are therefore not specific to either HA or NA. To resolve antigen specificity, we measured IgG responses against recombinant HA and NA from H1N1 and H5N1. Vaccinated males and females mounted comparable high anti-HA IgG titers against homologous HA antigens, specifically H1 in H1N1 vaccinated mice (**Fig. 1C**) and H5 in H5N1 vaccinated mice (**Fig. 1D**). Cross-reactive HA-specific IgG responses to heterologous viruses were low in both sexes and did not differ between males and females (**Fig. 1C-D**). NA-specific IgG titers against homologous N1 antigen were greater in females than males vaccinated against either H1N1 or H5N1 (**Fig. 1E-F**). H1N1 vaccinated females had greater NA-specific IgG responses to heterologous N1 antigen than males, whereas no sex differences were detected in heterologous anti-N1 responses in H5N1 vaccinated mice (**Fig. 1E-F**). Females mounted stronger virus-specific IgG responses than males, with stronger evidence of sex differences in cross-reactive total IgG and anti-NA IgG responses in H1N1 vaccinated mice.

**Fig. 1.**
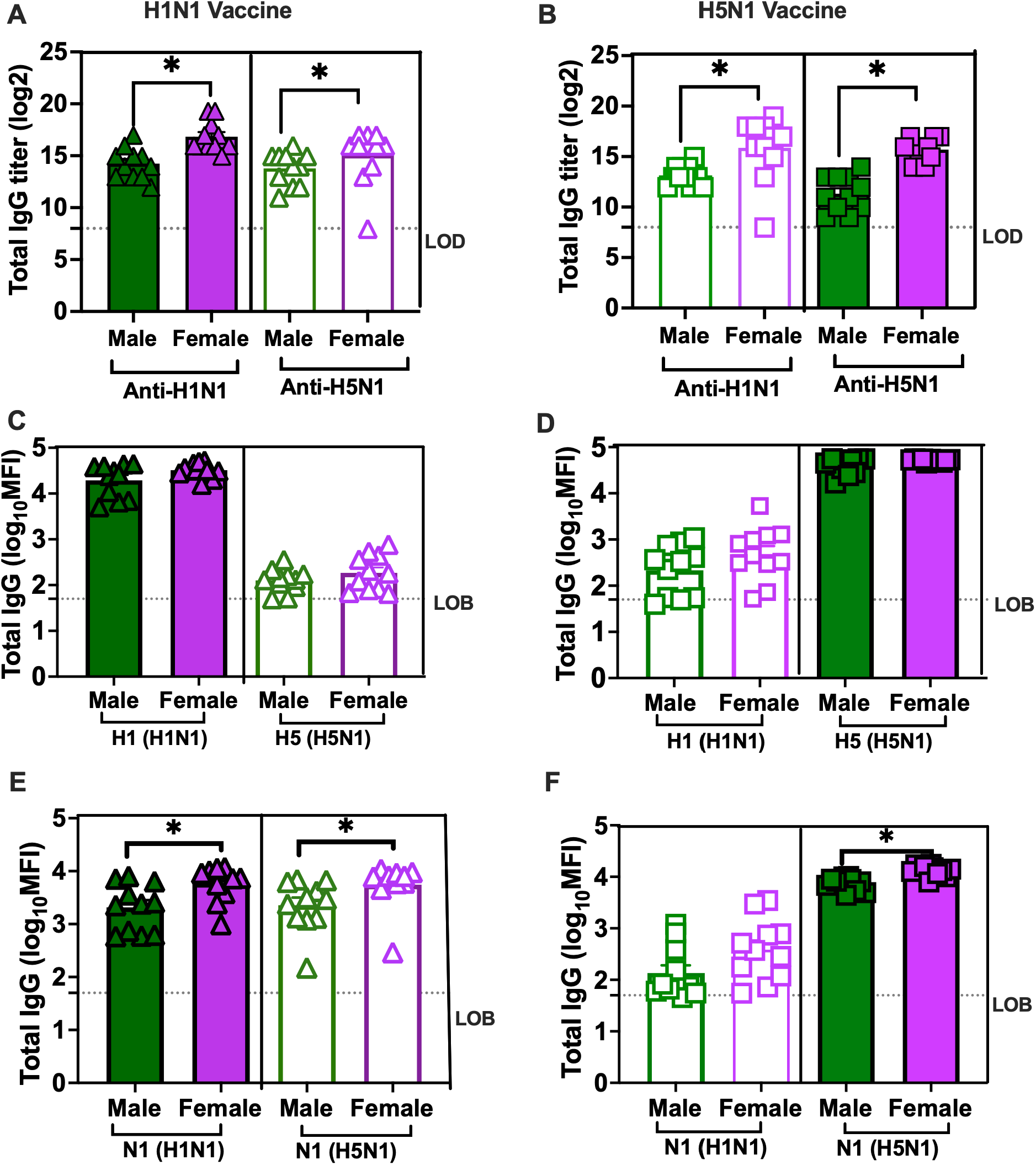
Females have greater homologous and heterologous virus-specific IgG titers than males. Adult (7–8-week-old) male (dark green) and female (dark purple) C57BL/6 mice were vaccinated twice with BPL-inactivated whole 2009 H1N1 or H5N1 (LAIV backbone) viruses using a prime/boost regimen with a 3-week interval. Plasma samples were collected 35 days post-vaccination to determine total anti-2009 H1N1 and anti-H5N1 IgG titers through ELISA in mice vaccinated with H1N1 (A) or H5N1 (B) vaccines. ELISA used whole virus coating antigen. Homologous and cross-reactive HA- and NA-specific total IgG responses were measured by multiplex systems serology and are shown as log10 mean fluorescence intensity (MFI) values for H1N1 HA (C), H5N1 HA (D), H1N1 NA (E), and H5N1 NA (F). Data represent mean ± SEM (n = 11/group). Asterisks (*) indicate significant sex differences (p < 0.05) determined by Mann-Whitney tests, using GraphPad Prism 10.1.0. LOB = limit of background; LOD = limit of detection.

### Females have greater homologous and heterologous NA-specific IgG2b responses with enhanced FcγRIV binding than males

Because females had greater homologous and heterologous IgG titers than males, we next determined which IgG subclass was contributing to the sex differences. In mice, total IgG consists of IgG1, IgG2b, IgG2c/a, and IgG3. Recent studies suggest that anti-H1N1 IgG2b is elevated in vaccinated female compared with male mice, which is caused by elevated concentrations of estradiol (22). We also previously reported greater H1N1-specific IgG2c in females than males following IIV (12). Following either H1N1 or H5N1 vaccination, IgG2b (**Fig 2A-D**) and IgG2c (**Fig. S1I-L**) were the most highly detectable isotypes produced in response to homologous and heterologous HA and NA. IgG1 and IgG3 titers were considerably lower than IgG2b/c responses against homologous HA and were undetectable against homologous NA and heterologous HA and NA following H1N1 vaccination (**Fig. S1M-T**). In contrast, H5N1 vaccination resulted in IgG1 titers that were within in same range as IgG2b/c against homologous HA and NA but undetectable against heterologous antigens (**Fig. S1M-P**). IgG3 titers following H5N1 vaccination were robust against homologous HA but frequently at or below the limit of background against homologous NA and heterologous antigens (**Fig. S1Q-T**).

**Fig. 2.**
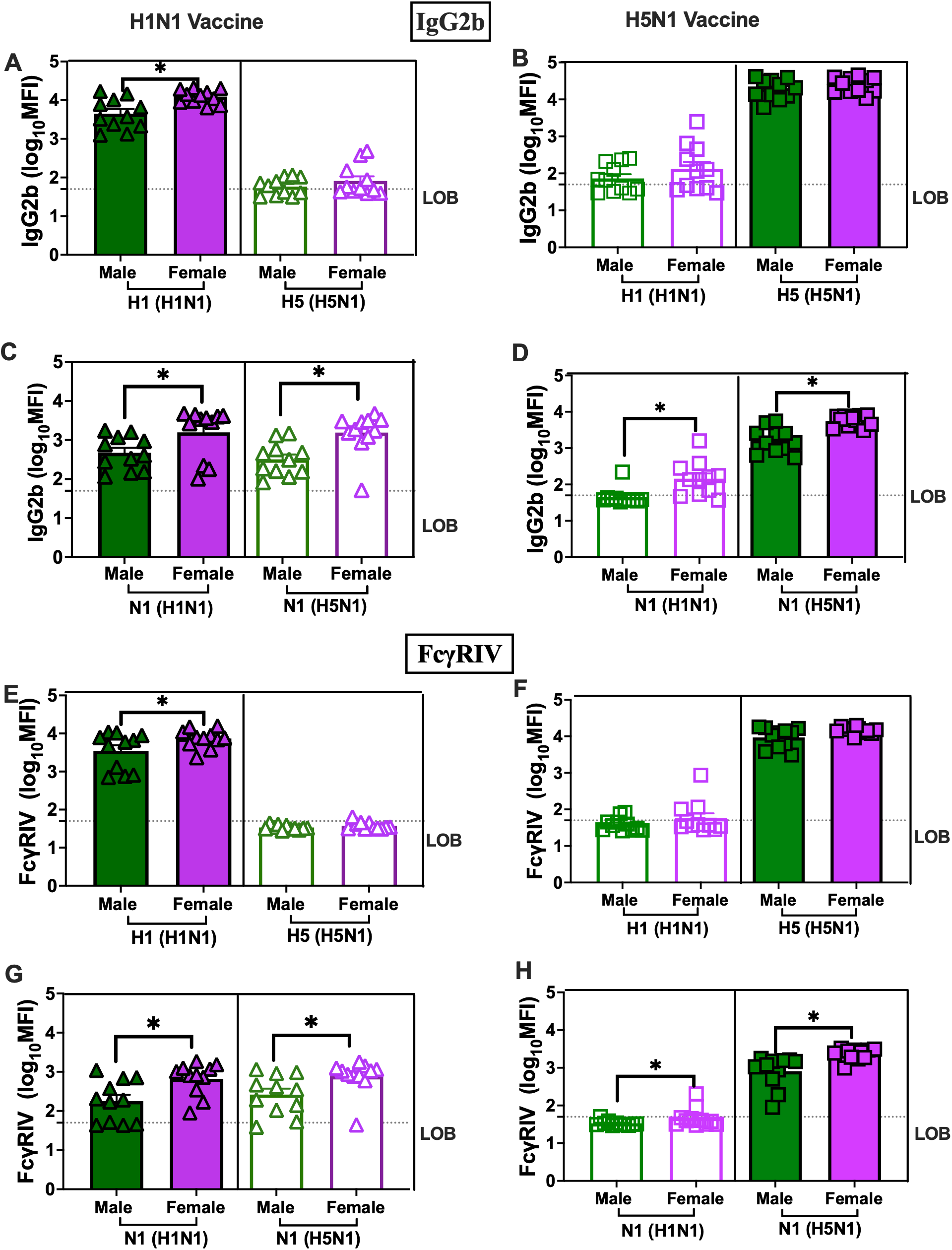
Females have greater homologous and heterologous NA-specific IgG2b responses with enhanced FcγRIV binding than males. Adult (7–8-week-old) male (dark green) and female (dark purple) C57BL/6 mice were vaccinated twice with BPL-inactivated whole 2009 H1N1 or H5N1 (LAIV backbone) viruses using a prime/boost regimen with a 3-week interval. Plasma samples were collected 35 days post-vaccination to determine homologous and cross-reactive HA- and NA-specific IgG2b responses by multiplex systems serology and are shown as log10 mean fluorescence intensity (MFI) values for H1 HA (A), H5 HA (B), H1N1 N1 (C), and H5N1 N1 (D). FcγRIV binding to antigen-specific antibodies was also measured by multiplex systems serology and is shown as log10 MFI values for H1 HA (E), H5 HA (F), H1N1 N1 (G), and H5N1 N1 (H). Data represent mean ± SEM (n = 11/group). Asterisks (*) indicate significant sex differences (p < 0.05) determined by Mann-Whitney tests, using GraphPad Prism 10.1.0. LOB = limit of background.

Sex differences were only observed in titers of IgG2 subclasses. H1N1 vaccinated females had greater IgG2b titers against homologous HA and NA, and significantly greater IgG2b titers against heterologous NA, but not HA, than their male counterparts (**Fig. 2A** and **2C**). H5N1-vaccinated females had greater IgG2b titers against homologous and heterologous NA, but not HA, than H5N1-vaccinated males (**Fig. 2B** and **2D**). A similar pattern was observed for IgG2c in which H1N1 vaccinated females had greater titers than males against both homologous and heterologous N1, and H5N1 vaccinated females had greater titers than males against homologous N1 (**Fig. S1I-L**). Overall, cross-reactive N1-specific IgG2 responses appeared to drive sex differences in total IgG responses (**Fig. 1**).

IgG isotypes have differential binding to Fcγ receptors (FcγR) I-IV, with mouse IgG2b preferentially engaging FcγRIV (23). Anti-HA and anti-NA antibodies from H1N1 vaccinated females had greater binding affinity to FcγRIV in response to homologous proteins than did antibodies from H1N1-vaccinated males (**Fig. 2E** and **2G**). Anti-NA antibodies from H1N1 vaccinated females showed greater binding to FcγRIV than those from males against heterologous NA (**Fig. 2G**). Anti-HA and anti-NA antibodies from H5N1 vaccinated mice exhibited stronger FcγRIV binding to homologous HA and NA, respectively, with limited cross-reactivity to heterologous proteins (**Fig. 2F** and **2H**). Anti-NA antibodies from H5N1 vaccinated females showed greater binding affinity to FcγRIV in response to homologous and heterologous antigens (**Fig. 2H**). Overall, FcγRIV binding was greater for vaccinated females than males in response to homologous and heterologous proteins. To determine whether greater binding to FcγRIV was a reflection of greater affinity to FcγRIV or greater amounts of IgG in females, homologous NA specific IgG: FcγRIV ratios were measured and were greater in males than females following either H1N1 or H5N1 vaccination, as were ratios of responses to homologous HA (**Fig. S2A-D)**. A similar trend was observed for heterologous N1 in H1N1 vaccinated mice, although this did not reach significance (**Fig. S2C**). These data illustrate that sex differences in FcγRIV engagement by antibodies recognizing homologous antigens is not explained by antibody quantity alone. The antibody binding to FcγRI, FcγRII, and FcγRIII was less robust than to FcγRIV in response to homologous and heterologous HA or NA (**Fig. S3A-L**). FcγRI-binding by homologous and heterologous HA- and NA-targeting antibody was comparable in both vaccine groups, with greater NA-specific homologous and cross-reactive binding observed in H5N1-vaccinated males compared to females (**Fig. S3C-D**). FcγRII and FcγRIII binding remained at or below background levels in response to homologous and heterologous antigens in H1N1 and H5N1 vaccinated mice (**Fig. S3E-L**). Together, these findings illustrate that vaccinated females produce greater IgG2 titers against NA from homologous and heterologous viruses, which corresponds with greater FcγRIV engagement, particularly following H1N1 vaccination.

### Functional antibody responses against homologous and heterologous viruses are greater in vaccinated females than males

Antibody-dependent complement deposition (ADCD) was evaluated as an Fc-mediated effector function. Robust homologous and heterologous HA and NA-specific ADCD was detected in sera from both H1N1- (**Fig. 3A** and **3C**) and H5N1-vaccinated (**Fig. 3B** and **3D**) mice. H1N1 and H5N1 vaccinated females had greater ADCD against homologous, but not heterologous, HA antigens than males (**Fig. 3A-B**). There were no sex differences in ADCD responses to either homologous or heterologous N1 among H1N1 vaccinated mice (**Fig. 3C**). H5N1 vaccinated males had greater ADCD against heterologous, but not homologous, N1 antigen than females (**Fig. 3D**). These findings suggest vaccination with inactivated H1N1 and H5N1 induces non-neutralizing ADCD antibody, with more consistent sex differences observed in response to homologous HA.

**Fig. 3.**
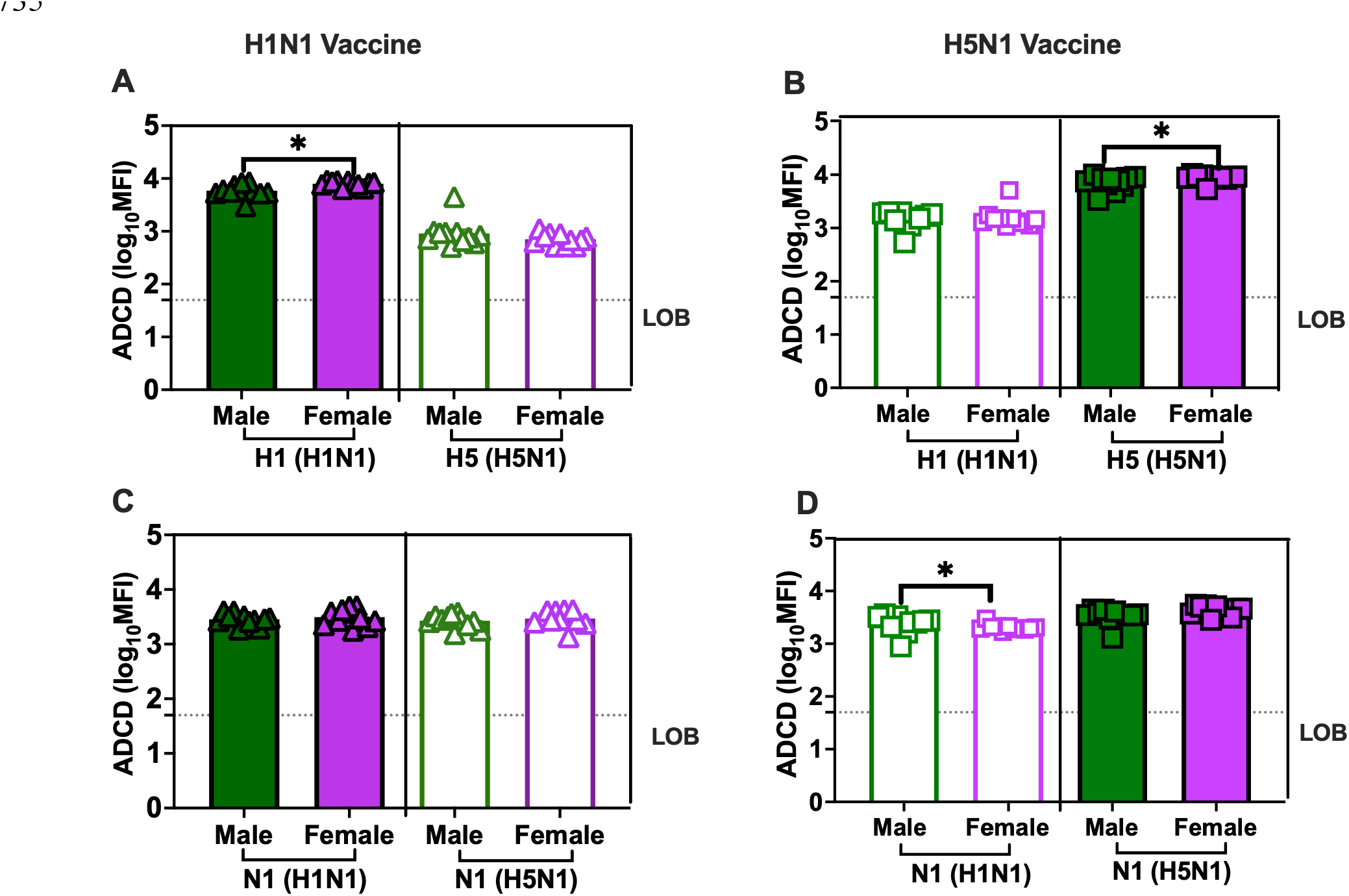
Antibody-dependent complement deposition (ADCD) responses are broadly induced following H1N1 and H5N1, with greater responses against homologous HA in females. Adult (7–8-week-old) male (dark green) and female (dark purple) C57BL/6 mice were vaccinated twice with BPL-inactivated whole 2009 H1N1 or H5N1 (LAIV backbone) viruses using a prime/boost regimen with a 3-week interval. Plasma samples were collected 35 days post-vaccination to assess HA- and NA-specific antibody-dependent complement deposition (ADCD). Biotinylated antigen-coated beads were incubated with diluted plasma (1:40) to allow immune complex formation, followed by measurement of C3b deposition as a readout of complement activation. ADCD responses are shown as mean fluorescence intensity (MFI) values for H1 HA (A), H5 HA (B), H1N1 N1 (C), and H5N1 N1 (D). Data represent mean ± SEM (n = 10–11/group). Asterisks (*) indicate significant sex differences (p < 0.05) determined by Mann-Whitney tests, using GraphPad Prism 10.1.0. LOB = limit of background.

Vaccinated female mice tended to generate greater homologous nAb titers than males post-vaccination (p=0.07) (**Fig. 4A-B**). No cross-reactive nAbs were detected in either vaccine group (**Fig. 4A-B**). Because nAb against influenza viruses are predominantly mediated by HA-specific antibody, these data highlight that cross-reactive HA antibody following vaccination are likely non-neutralizing in activity.

**Fig. 4.**
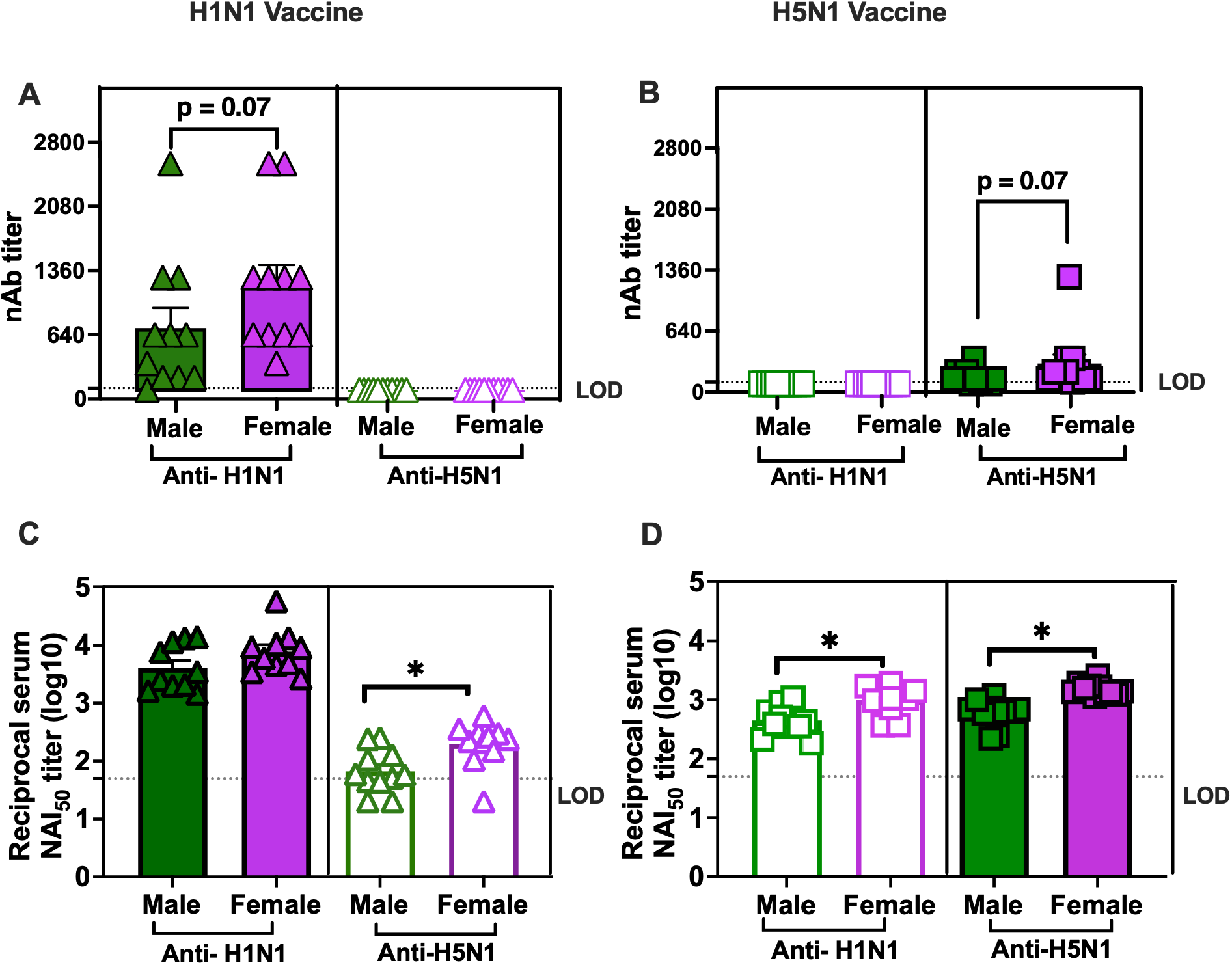
Females develop stronger homologous and heterologous functional antibody responses following H1N1 and H5N1 vaccination. Adult (7–8-week-old) male (dark green) and female (dark purple) C57BL/6 mice were vaccinated twice with BPL-inactivated whole 2009 H1N1 or H5N1 (LAIV backbone) viruses using a prime/boost regimen with a 3-week interval. Plasma samples were collected 35 days post-vaccination to determine homologous and heterologous virus-neutralizing antibody (nAb) titers against 2009 H1N1 and H5N1 (LAIV) viruses in mice vaccinated with H1N1 (A) or H5N1 (B) vaccines. Neuraminidase inhibition (NAI) antibody responses against H5N1(LAIV) and H1N1 viruses were measured by enzyme-linked lectin assay in mice vaccinated with H1N1 (C) or H5N1 (D) vaccines. Data represent mean ± SEM (n = 10–11/group). Asterisks (*) indicate significant sex differences (p < 0.05) determined by Mann-Whitney tests, using GraphPad Prism 10.1.0. LOB = limit of background.

NA-specific antibodies that inhibit NA enzymatic activity are associated with heterologous protection against H5N1 in both ferret and mouse models (24, 25). Because heterologous NA binding antibodies in vaccinated mice were greater in females than males, we aimed to determine whether NA enzyme activity inhibition by antibodies (i.e., NAI antibodies) from either H1N1 or H5N1 vaccinated mice could be achieved. In H1N1-vaccinated mice, while both males and females had equally high homologous NAI titers against H1N1, females had greater heterologous NAI titers against H5N1 than males (**Fig. 4C**). In H5N1-vaccinated mice, females had greater homologous and heterologous NAI titers than males (**Fig. 4D**). Overall, NAI titers were greater in vaccinated females than males.

### H1N1 vaccination enables females but not males to limit H5N1 virus dissemination

Naïve adult male and female mice were infected with one of two human-origin H5N1 strains, A/Texas/37/2024 or A/Michigan/90/2024 at equivalent doses and followed them for morbidity and mortality (**Fig. S4A-F**). Infection with A/Texas/37/2024 resulted in more rapid body mass loss and development of clinical disease (**Fig. S4A** and **4C**) and earlier mortality (**Fig. S4E**) as compared with A/Michigan/90/2024 (**Fig. S4B, 4D**, and **4F**). In response to A/Texas/37/2024, survival was significantly lower for females than males, with females dying 2 days earlier than males (**Fig. S4E**). In contrast, A/Michigan/90/2024 infection resulted in comparatively attenuated disease, with no apparent sex differences in survival (**Fig. S4F**), despite observable sex differences in body mass loss and clinical severity scores (**Fig. S4B** and **4D**).

To assess whether BPL-inactivated H1N1 and H5N1 vaccines could protect mice against H5N1 infection, adult males and females were either mock vaccinated or vaccinated with H1N1 or H5N1 and challenged with a lethal dose of A/Texas/37/2024 at 48 dpv. Because the H5N1 vaccine contains the HA and NA of a clade 2.3.4.4b virus closely related to the challenge virus, this group represented a homologous vaccine challenge control, whereas the H1N1 vaccinated group, with heterologous HA and NA, was the heterologous comparison. Both H1N1 and H5N1 vaccination conferred 100% survival in both males and females through 5 dpi, with H1N1- and H5N1-vaccinated mice maintaining greater body mass than mock-vaccinated mice (**Fig. 5A**). Clinical scores remained low in vaccinated males and females, with no sex differences observed (**Fig. S5A-B**). Virus replication in nasal turbinate (**Fig. 5B**), trachea (**Fig. 5C**), and lung (**Fig. 5D**) tissue was quantified to evaluate how well vaccine-induced immunity could limit virus replication. H5N1 vaccination resulted in reduced virus titers as compared with mock-vaccinated mice in the tracheas and lungs, with no sex differences observed at 3 days post challenge (dpc) (**Fig. 5B-D**). At 3 dpc, very little replicating virus was detected in nasal turbinates, with low H5N1 titers detectable in n=2 mock vaccinated males and n=1 H5N1 vaccinated females (**Fig. 5B**). In the tracheas (i.e., upper respiratory tract), H1N1 and H5N1 vaccination reduced virus titers as compared with mock vaccination (**Fig. 5C**). Among H1N1-vaccinated mice, both males (3/5) and females (4/6) had detectable H5N1 virus in tracheas (**Fig. 5C**), with no sex differences observed. In the lungs (i.e., lower respiratory tract), H5N1 vaccination resulted in non-detectable virus in all but one female mouse. Among H1N1 vaccinated mice, females had significantly lower virus titers than males at 3 dpc (**Fig. 5D**), with both H1N1 and H5N1 vaccinated females having lower pulmonary virus titers than mock vaccinated females (**Fig. 5D**). To determine whether virus disseminated beyond the respiratory tract, gonad tissues and seminal fluid were analyzed for detectable virus at 3 dpc. While infectious virus was detected in 2/5 ovaries from mock-vaccinated females, no virus was detected in ovaries from vaccinated females (**Fig. S5C**). Live virus also was not detected in testes or seminal fluid from mock-, H1N1-, or H5N1-vaccinated males (**Fig. S5D-E**) suggesting that at least at this early timepoint, virus did not disseminate to reproductive tissues. These data highlight that H1N1 vaccination enables females, but not males, to restrict virus replication to the upper respiratory tract, which may involve greater induction of heterologous NAI antibody responses in H1N1 vaccinated female mice.

**Fig. 5.**
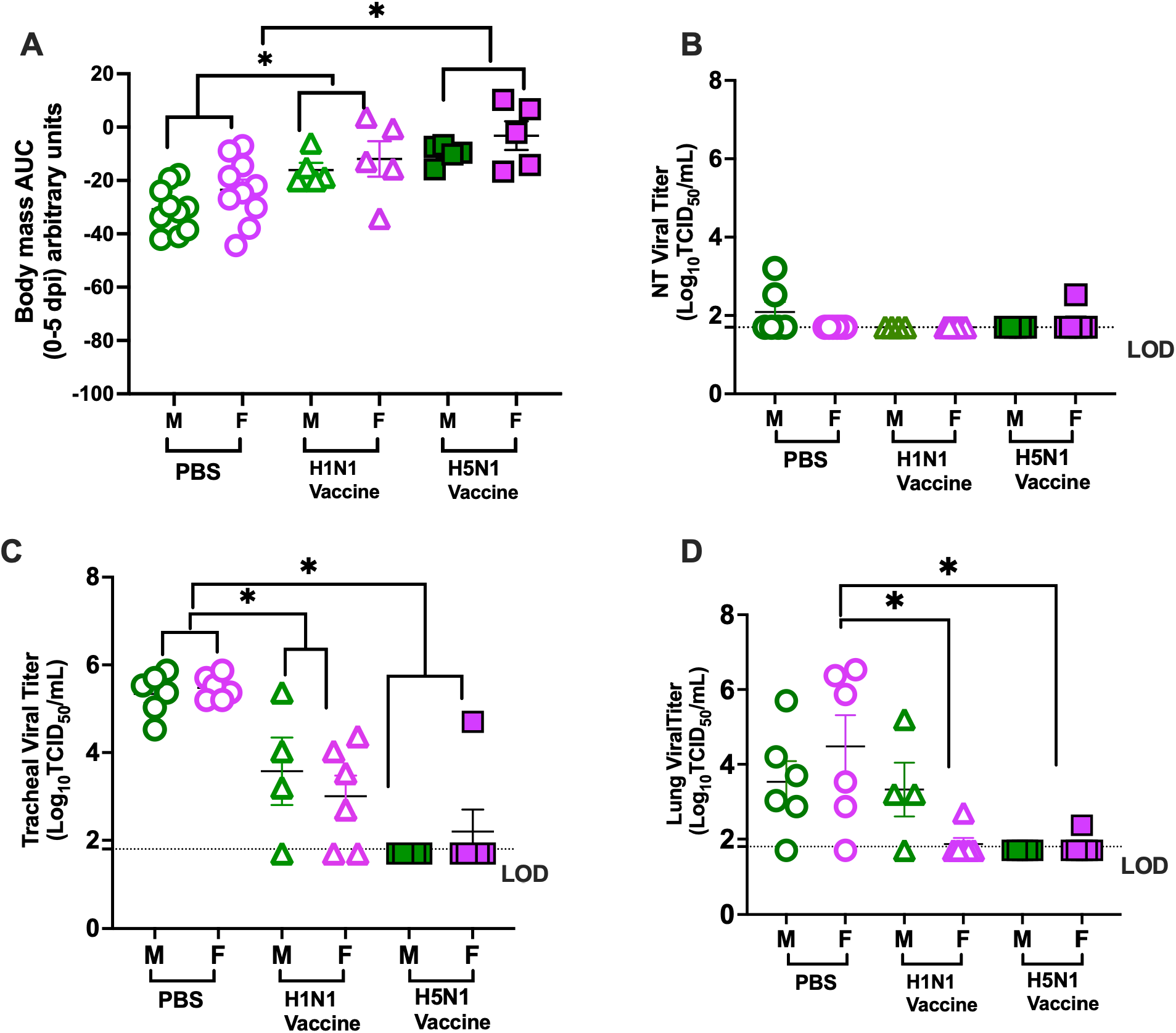
H1N1 vaccination provides greater control of H5N1 respiratory viral replication in females than males. Adult 7–8-week-old male (dark green) and female (dark purple) C57BL/6 mice were mock vaccinated with PBS or vaccinated twice, with a 3-week interval, using BPL-inactivated whole 2009 H1N1 or H5N1 (LAIV backbone) viruses in a prime/boost regimen. Six weeks after vaccination, mice were challenged with 10^2^ TCID_50_ of A/human/Texas/37/2024 (H5N1). To assess morbidity, percent change in body mass was measured daily for 5 days post-challenge (mock-vaccinated, n = 10/sex; H1N1-vaccinated, n = 5-6/sex; H5N1-vaccinated, n = 5/sex). Virus titers in the nasal turbinate (NT) (B), trachea (C), and lung (D) were measured on day 3 post-challenge by TCID_50_ assay (n = 5–6/group). Data are presented as mean ± SEM. Asterisks (*) indicate significant differences between PBS mock-vaccinated and vaccinated groups (p < 0.05) as determined by two-way ANOVA using GraphPad Prism 10.1.0. LOD = limit of detection.

## Discussion

Biological sex shapes influenza vaccine-induced cross-recognizing immunity and virus dissemination following H5N1 infection. Female mice vaccinated against either H1N1 or H5N1 mounted greater virus-specific IgG and homologous neutralizing antibody responses than males. Vaccinated females also exhibited greater cross-reactive immunity against heterologous viruses. Cross-reactive immunity was predominantly observed in response to heterologous NA, rather than HA, antigens. Females also exhibiting greater NAI titers against heterologous viruses, particularly among mice that received H1N1 vaccine. Following live H5N1 challenge, H1N1 vaccinated females had lower virus titers in their lungs than males, whereas males and females that received the H5N1 vaccine were equally protected. These findings suggest that cross-reactive immunity against H5N1 is mediated by antibody responses to NA, which might contribute to greater restriction of H5N1 challenge virus in the upper respiratory tract of females compared with males. Our use of whole, inactivated virus as a vaccine likely contributes to robust NA-specific responses, as conventional inactivated influenza vaccines (i.e., split or subunit) frequently contain reduced or antigenically compromised NA (16, 26, 27).

We multiplexed a broad panel of IgG isotypes and binding affinity to FcγRs. IgG isotypes differ in their ability to mediate Fc-dependent effector functions (23, 28). In mice, IgG2 subclasses generally exhibit stronger Fc-mediated antiviral activity due to their broader binding affinity FcγRI, FcγRII, FcγRIII, and FcγRIV (28). IgG2 in mice, which is similar to IgG1 in humans, is considered a particularly important antiviral subclass because it efficiently activates complement and engages FcγRIV on monocytes, macrophages, neutrophils, and other inflammatory myeloid cells, thereby promoting phagocytosis, immune complex clearance, cytokine production, and antibody-dependent cellular cytotoxicity-like activity (23). Elevated IgG2 responses in mice have been associated with improved viral clearance and enhanced heterologous protection following influenza infection and vaccination, particularly when neutralizing antibody responses are limited, with cross-reactive non-neutralizing IgG2 mediating Fc-dependent protection against heterologous viral challenge (29, 30). In our study, vaccinated females exhibited greater homologous and cross-reactive IgG2 responses against HA and NA than males, suggesting enhanced Fc-mediated antiviral immunity in females. Together, these findings suggest that sex differences in IgG2 responses may underlie the enhanced breadth of influenza vaccine-induced protection in females, highlighting IgG2-dominant responses as a potential target for improving vaccine breadth and effectiveness.

Non-neutralizing, complement-activating antibodies can contribute to influenza immunity by promoting membrane attack complex formation, leading to lysis and clearance of virus-infected targets. Influenza vaccination induces ADCD as part of a broader Fc-effector mechanism and complement component C3 has been shown to be essential for non-neutralizing antibody-mediated protection against influenza (31–35). Both HA- and NA-specific antibodies mediate broadly cross-reactive and comparable ADCD responses between males and females, indicating a robust Fc-effector function that may contribute to some heterologous protection.

Females showed greater ADCD against homologous HA in both H1N1 and H5N1 vaccine groups, whereas no consistent sex differences were observed against the N1 from either H1N1 or H5N1. Additional Fc-effector mechanisms include FcγR–mediated phagocytosis and cytotoxicity, which also may enhance macrophage and NK cell-mediated cross-protection through clearance of infected cells (36–38), was not investigated in the current study.

Influenza virus NA promotes replication by cleaving terminal sialic acid residues to facilitate viral release and dissemination from infected cells and may also contribute to early stages of infection by enhancing viral entry into host epithelial cells (1, 2). NA-specific antibodies that inhibit neuraminidase activity can limit viral replication by blocking virus release and cell-to-cell spread and have been associated with reduced disease severity and decreased viral shedding following influenza infection (3, 39). Cross-reactive NAI responses are associated with reduced disease severity and improved survival against heterologous influenza virus infection (24, 40, 41). Structural analyses show that the N1 of pandemic H1N1 viruses share greater sequence conservation with avian H5N1 than with seasonal human H1N1 strains, providing a structural basis for cross-recognition (8). Notably, the magnitude of cross-reactive N1 binding and NA inhibition were not directly proportional. Binding assays capture antibodies recognizing multiple regions of NA protein, whereas inhibition depends on whether antibody binding interferes with substrate access to the enzymatic site (42). Thus, H1N1 vaccination may generate a broader cross-reactive N1 antibody response, resulting in greater cross-reactive N1 binding but comparatively lower inhibitory antibodies. Beyond NAI inhibition, these antibodies also might contribute to antiviral immunity through other antibody-mediated mechanisms, including Fc-dependent effector functions (43). In the current study, vaccinated females mounted stronger heterologous NA-specific antibody responses and H1N1-vaccinated females had greater NAI titers than their male counterparts– which was associated with reduced H5N1 titers after live virus challenge. These data suggest that sex differences in NA-mediated immunity affect heterologous influenza protection, highlighting the importance of considering biological sex in the design and evaluation of next-generation influenza vaccines aimed at enhancing NA-mediated cross-protective immunity.

This study has several limitations. First, we evaluated immune responses using only whole inactivated influenza vaccines, and whether similar sex differences in NA-specific cross-reactive immunity occur with other vaccine platforms, including live-attenuated, recombinant protein, mRNA, or viral vectored vaccines, remains to be determined. Second, our studies were performed in immunologically naive mice, which do not recapitulate the complex pre-existing influenza immunity present in humans resulting from prior infection and vaccination. Pre-existing antibodies and immune imprinting may substantially influence the magnitude, specificity, and breadth of cross-reactive responses (44, 45). Third, our analyses primarily focused on binding and functional antibody responses against the major surface glycoproteins HA and NA, whereas cellular immune responses to internal viral proteins were not evaluated (46)(47, 48). Because NP-specific and other internal protein-specific T cell responses can contribute to heterologous influenza protection, future studies should investigate how biological sex influences cross-reactive T cell responses following influenza vaccination and infection (49–51). Although inactivated influenza vaccines generally induce limited T cell immunity, defining the contribution of viral antigen specific cellular responses alongside antibody responses will be important for understanding the broader mechanisms underlying sex differences in heterologous influenza protection (11, 52). Finally, virus titers only were assessed at 3 dpc which did not capture the kinetics of virus clearance.

In conclusion, our study demonstrates that H1N1 vaccinated females develop stronger NA-directed functional and cross-reactive immune responses than males following influenza vaccination, resulting in enhanced virus control following heterologous H5N1 challenge. These findings identify NA-specific immunity as a major contributor to limiting heterologous virus dissemination and support NA as an important target for next-generation influenza vaccines.

Future human studies should consider using adjuvants and vaccine platforms that enhance NA-directed and Fc-mediated immune responses that may broaden the heterosubtypic immune responses and protection. Human studies of NA-mediated immunity must consider that biological sex could impact outcomes.

## Materials and methods

### Mice

Adult (7–8-week-old) male and female C57BL/6 mice (n = 5-11/group) were obtained from Jackson Laboratories. Mice were housed five per cage under standard animal biosafety level (ABSL)-2 conditions, with *ad libitum* food and water throughout the vaccination regimen. For live virus challenge experiments, mice were transferred to standard ABSL-3 animal facilities for H5N1 infection. Mice were given more than 24 h to acclimate the ABSL3 facility before infections (53). All work with H5N1 virus was approved and performed in accordance with biosafety and operating procedures approved by the Johns Hopkins University Biosafety Office (registration #: P2406130101). All animal procedures were approved by the Johns Hopkins University Animal Care and Use Committee (Protocol MO24H158) and the Johns Hopkins Biosafety Office.

### Cells

Madin-Darby canine kidney (MDCK) cells, a kind gift from Dr. Robert A. Lamb, were maintained in complete medium (CM) consisting of Dulbecco’s Modified Eagle Medium (DMEM, Sigma-Aldrich, St Louis, MO, USA) supplemented with 10% fetal bovine serum (FBS; Gibco, Waltham, MA, USA), 100 units/ml penicillin/streptomycin (Life Technologies, Frederick, MD, USA) and 2 mM Glutamax (Gibco, Waltham, MA, USA) at 37 °C and 5% CO2.

### Vaccines and viruses

A/human/Texas/37/2024 (H5N1; A/human/TX) and A/human/Michigan/90/2024 (H5N1; A/human/Michigan) were kindly provided by the U.S. Centers for Disease Control. Virus stocks were generated by infecting MDCK cells at a multiplicity of infection (MOI) of 0.01 at 33°C in infection media (Complete media without FBS with 5 μg/ml of N-acetyl Trypsin and 0.3% bovine serum albumin (Sigma)) as previously described (7). A/California/04/09 (2009 H1N1) was generated by reverse genetics from a published sequence (20). For H5N1 LAIV virus vaccine, HA and NA segments from A/Bovine/Texas/24-029328-01/2024 (GISAID accession no. EPI_ISL_19014384) were cloned into an eight-plasmid expression system with bidirectional promoters as previously described (kindly provided by Andrew Bowman and Richard Webby) (7, 54, 55). The H5 HA-encoding plasmid had the multibasic cleavage site (MBS) deleted, to be in accordance with standard BSL-2 containment practices. The remaining six internal gene segments from A/Ann Arbor/6/1960 (H2N2) strain (GISAID accession no. EPI_ISL_130415), a cold-adapted virus (live attenuated, indicated as LAIV), were cloned into the 12-plasmid reverse genetics system and co-transfected with the bovine-derived A/H5N1 HA and NA into co-cultured MDCK and HEK-293T cells, as previously described (7, 54, 55).Viruses were inactivated with 0.05% (v/v) β-propiolactone (Millipore Sigma) prior to purification by ultracentrifugation with a sucrose cushion as previously described (11).

### Vaccination and H5N1 challenge

Mice were mock vaccinated with 1xPBS or immunized with 20 µg of beta-propiolactone inactivated whole virus A/California/04/2009 H1N1 (2009 H1N1) or H5N1 (LAIV backbone) vaccines, using a prime/boost strategy on a 3-week interval, administered intramuscularly in the right thigh muscle (11). Blood samples were collected into heparinized tubes at 35 days post-prime vaccination (dpv) through the retro-orbital route under isoflurane anesthesia. Three weeks after the final boost (48 dpv), under ketamine-xylazine anesthesia, vaccinated mice were intranasally challenged with 10^2^ TCID_50_ of the A/human/Texas/37/2024 (H5N1). Subsets of mice were humanely euthanized at 3 days post challenge (dpc), with changes in body mass and clinical disease recorded daily for 5 dpc, depending on the virulence of the virus. For survival, mice were either found dead or were humanely euthanized when body mass dropped below 20% starting mass. Mice were anesthetized using a ketamine-xylazine cocktail before intracardial bleeding. Blood was collected into heparinized tubes, and the plasma was separated by centrifugation at 9600 × g for 30 min at 4°C. Plasma was stored at −80°C and used to measure titers of antibodies. Nasal turbinate, tracheas, lungs, ovaries, testes and seminal fluid\ were collected, snap-frozen, and stored at −80 °C until processed for downstream analysis.

### Enzyme-linked Immunosorbent Assays (ELISAs)

Anti-H1N1 and anti-H5N1 IgG antibodies in plasma samples collected 35 dpv, using our in-house validated ELISAs (4, 12, 13). Briefly, ELISA plates were coated with 50 μL/well of PBS buffer containing 2µg/mL of BPL inactivated 2009 H1N1 or H5N1 LAIV whole virus by overnight incubation at 4 °C. Plates were washed three times, blocked for 1 h at 37 °C using 10% skim milk solution, and then two-fold serially diluted plasma samples were added. Plates were incubated at 37°C for 1 h and washed three times. Horseradish peroxidase (HRP)-conjugated secondary IgG antibodies (Invitrogen) were added and incubated for 1 h at room temperature. Plates were washed, and reactions were developed using 3,3’,5,5’-tetramethylbenzidine (TMB, BD Biosciences) for 20 min and stopped using 1 N hydrochloric acid (HCl). Using an ELISA plate reader (Molecular Devices), all plates were read at 450 nm wavelength, and the endpoint titer was calculated as the highest plasma dilution with an average optical density (OD) value greater than three times the average OD of mock (unvaccinated) animal sample as a negative control (56, 57).

### Virus titration

MDCK cells were cultured in CM. Cells were incubated in a 5% CO₂ humidified incubator at 37°C. For virus titration, frozen nasal turbinate, tracheas, lungs, gonads, and seminal fluids were homogenized. Homogenates were clarified by centrifugation at 12,000 rpm and stored at −80°C until analysis. Tissue homogenates were 10-fold serially diluted in serum-free media and transferred in six replicates into 96-well cell culture plates confluent with MDCK cells. Plates were incubated for 6 days at 32°C, followed by fixation with 4% formaldehyde, staining with naphthol blue-black solution, and virus titer calculation using the Reed-Muench method (58).

### Live virus neutralization

Heat-inactivated (56°C, 35 min) plasma samples were 2-fold serially diluted in infection medium (starting at a 1:20 dilution) and incubated with 100 TCID_50_ of H1N1 and H5N1 LAIV viruses. After a 1 h incubation at room temperature, the plasma-virus mix was transferred into a 96-well plate, confluent with MDCK cells in quadruplicate wells. The plasma and virus mixture was then used to infect quadruplicate wells of confluent MDCK cells for 24 h at 32°C. Following the 24 h incubation, the inoculum was removed and the cells were washed with 1X PBS and new media was added. The cells were incubated for 6 days at 32°C and then fixed with 4% formaldehyde and stained with naphthol blue black overnight. The neutralizing antibody titer was calculated as the highest plasma dilution that eliminated virus cytopathic effects in 50% of the wells (7, 11, 12).

### ELLA for quantification of NAI antibody response

ELLA was used to measure neuraminidase activity–inhibiting (NAI) antibodies, following a previously described method (7). Briefly, Nunc 96-well Immulon 4 HBX plates (Thermo Fisher) were coated with fetuin from FBS (Sigma) at 2.5 µg/well in 1× PBS, sealed, and incubated at 4°C for 12–16 h. Plasma samples were serially diluted fourfold for eight dilutions in assay buffer containing 0.2% Tween 20 (Sigma), 1% BSA (Sigma), 0.1 mg/mL MgCl₂, and 0.2 mg/mL CaCl₂ in PBS. A/H1N1 or A/H5N1-LAIV virus, diluted in the same buffer, was incubated with diluted sera at 37°C for 1 h. In parallel, fetuin-coated plates were washed 3× with PBST. Plasma–virus mixtures were added in duplicate, with control wells containing assay buffer only (0% NA activity/100% NAI) and virus only (100% NA activity/0% NAI). Plates were sealed and incubated for 16–18 h at 37°C with 5% CO₂. After washing with PBST, plates were incubated for 2 h at room temperature with HRP-conjugated peanut agglutinin (Arachis hypogaea lectin; Sigma). Following a final wash, SigmaFast OPD substrate (Sigma) was added for ∼10 min in the dark, the reaction was stopped with 1 N H₂SO₄, and absorbance was measured at 490 nm to determine relative NAI levels.

### Luminex Antigen-Bead Coupling

Luminex antigen–bead coupling was performed by covalently linking antigens to distinct MagPlex bead regions via a carboxylate activation reaction. Selected antigens included H1 HA (H1N1; Enzyme #IA-11SW-005P), H5 (H5N1; Sino Biological #41036-V08H), N1 from A/California/04/2009 and N1 from A/dairy cow/Texas/24-008749-002-v/2024 that were produced were provided by the Coughlan Lab. Beads were activated using activation buffer (0.1 M NaH₂PO₄, pH 6.2) with 50 mg/mL EDC and 50 mg/mL Sulfo-NHS (N-hydroxysulfosuccinimide). Following activation, 25 µg of antigen was coupled in coupling buffer (0.05 M MES, pH 5.0). Beads were then blocked with blocking buffer (PBS, 0.1% BSA, 0.02% Tween-20, 0.05% sodium azide; pH 7.4), washed with washing buffer (PBS, 0.05% Tween-20), and stored in PBS at 4°C until use.

### Multiplex Isotyping and FcγR Binding Assays

For all assays, 5 µL of plasma (1:40 dilution) was incubated with bead solution (20 µL per antigen-linked bead region in Luminex Assay Buffer: PBS 1X, 0.1% BSA, 0.05% Tween-20). Selected antigens include HA (H1 or H5) and NA (N1 from either H1N1 or H5N1). Incubation was performed for 2 h on a plate shaker, followed by magnetic washing using Procartaplex Wash Buffer (Thermo Fisher). Isotype and FcγR detection stains were prepared separately in conical tubes. For isotype analysis, PE-conjugated antibodies (Southern Biotech) were used to detect antigen-specific IgG (IgG1–4), IgM, and IgA. A total of 45 µL of isotype stain was added per well according to the assay layout. For FcγR binding analysis, FcγRI, FcγRII, FcγRIII, and FcγRIV proteins (Duke Vaccine Institute Protein Production Facility) were biotinylated at least 1 day prior using a BirA biotin-ligation kit (Avidity) and coupled to PE-streptavidin (BioLegend) for 15 min on a rotor before use. A total of 45 µL of FcγR probe stain was added per well according to the assay layout. Mean fluorescence intensity (MFI) of PE-conjugated secondary antibodies or FcγR probes was measured for each bead region using a Luminex Intelliflex system to quantify isotype levels and FcγR binding responses. Samples were acquired on a FlexMap 3D instrument (low PMT setting) with a doublet discrimination range of 7000–17000, and a minimum of 100 events per antigen. Assay background was defined using PBS as a negative control, and positive controls were included to ensure assay integrity.

### Antibody-Dependent Complement Deposition (ADCD) Assay

Biotinylated antigen-coated beads were incubated with diluted plasma (1:40) to allow immune complex formation. Selected antigens HA (H1 or H5) and NA (N1 from either H1N1 or H5N1). Lyophilized Low-Tox Guinea Pig Complement (Cedarlane) was reconstituted in ice-cold dH_2_O and diluted 1:25 in Gelatin Veronal Buffer with Ca^2+^ and Mg^2+^ (GVB++; Boston Bio). Diluted complement was added to each well and incubated at 37°C to allow complement activation and deposition. Complement binding was detected using goat anti-guinea pig C3 antibody (MP Biomedicals, 1:500 in 0.1% BSA), followed by donkey anti-goat IgG-PE secondary antibody (Thermo Fisher, 1:200 in 0.1% BSA). Beads were resuspended in 0.1% BSA and acquired on a Luminex Intelliflex system as described above.

### Statistical analyses

Data were analyzed using GraphPad Prism® version 10.1.0. Antibody measurements, including whole-virus and antigen-specific IgG titers determined by ELISA and multiplex bead-based assays, IgG subclass titers, FcγRIV binding, ADCD, neutralization titers, and neuraminidase inhibition (NAI) titers, were compared between males and females using unpaired Mann– Whitney tests. Area under the curve (AUC) measurements for body mass and tissue virus titers were compared using two-way ANOVAs followed by Tukey post hoc multiple-comparison tests. Morbidity data were analyzed using multivariate ANOVA. Survival distributions were compared using Kaplan–Meier analyses with the log-rank test. Statistical significance was defined as p < 0.05.

## Acknowledgements

We thank the expert animal care staff at the Miller Research Building Animal Facilities at Johns Hopkins University for assistance with animal studies Johns Hopkins University and the Klein, Davis, Thompson, and Pekosz lab members for discussions.

## Data availability

All data used in the study are available at https://doi.org/10.7281/T1RBFU3H

---

**Fig. S1.**
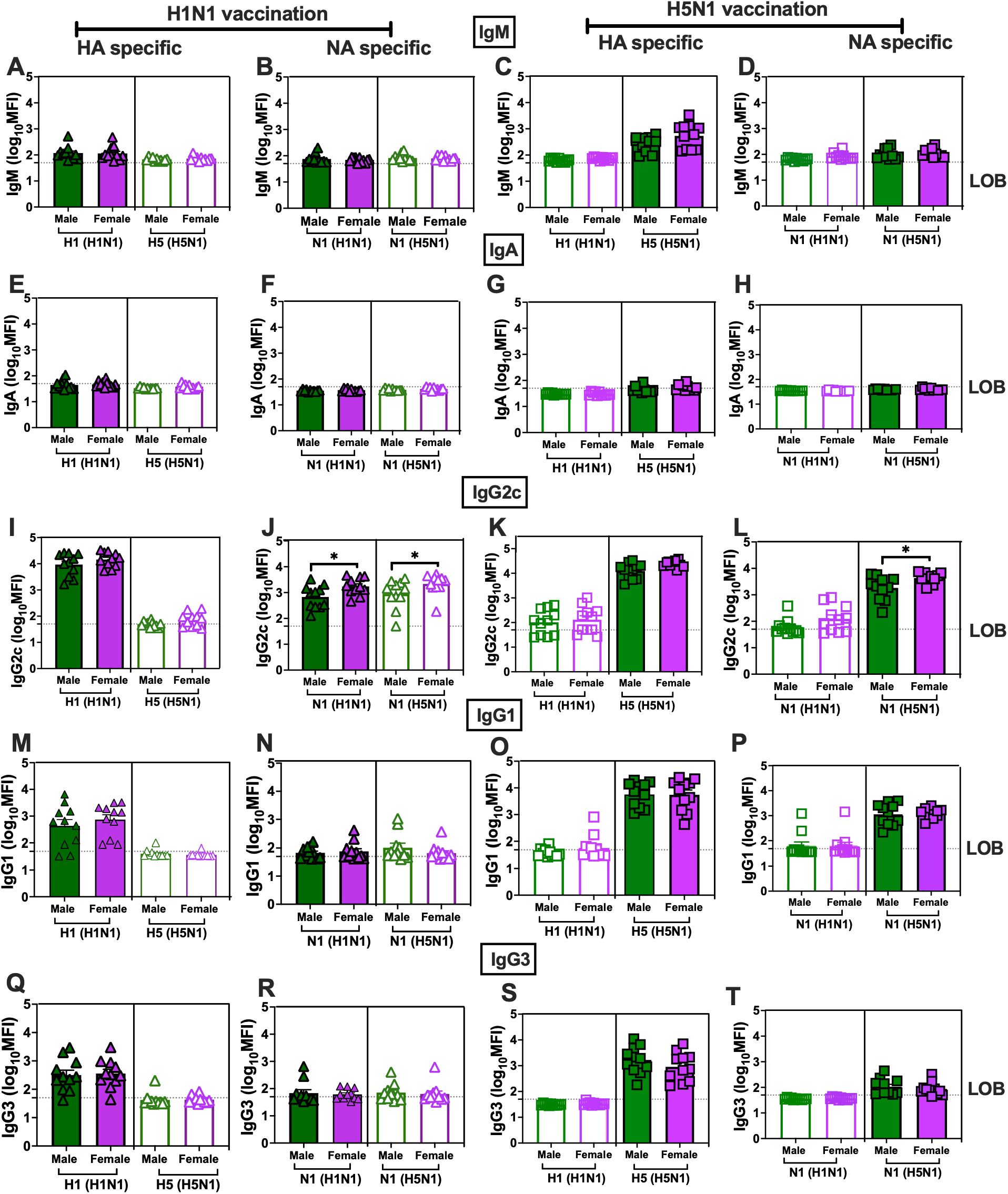
Homologous and cross-reactive HA- and NA-specific antibody responses following H1N1 and H5N1 vaccination in C57BL/6 mice. Adult 7–8-week-old male (dark green) and female (dark purple) C57BL/6 mice were vaccinated twice, with a 3-week interval, using BPL-inactivated whole 2009 H1N1 or H5N1 (LAIV backbone) viruses in a prime/boost regimen. Plasma samples were collected 35 days post-vaccination. Homologous and cross-reactive HA- and NA-specific IgM (A–D), IgA (E–H), IgG2c (I–L), IgG1 (M–P), and IgG3 (Q–T) responses were measured by multiplex systems serology and are shown as log10 mean fluorescence intensity (MFI) values for the selected antigens, HA (H1 or H5) and NA (N1 from either H1N1 or H5N1). Data are presented as mean ± SEM (n = 10–11/group). Asterisks (*) indicate significant sex differences (p < 0.05) determined by Mann-Whitney tests, using GraphPad Prism 10.1.0. LOB = limit of background.

**Fig. S2.**
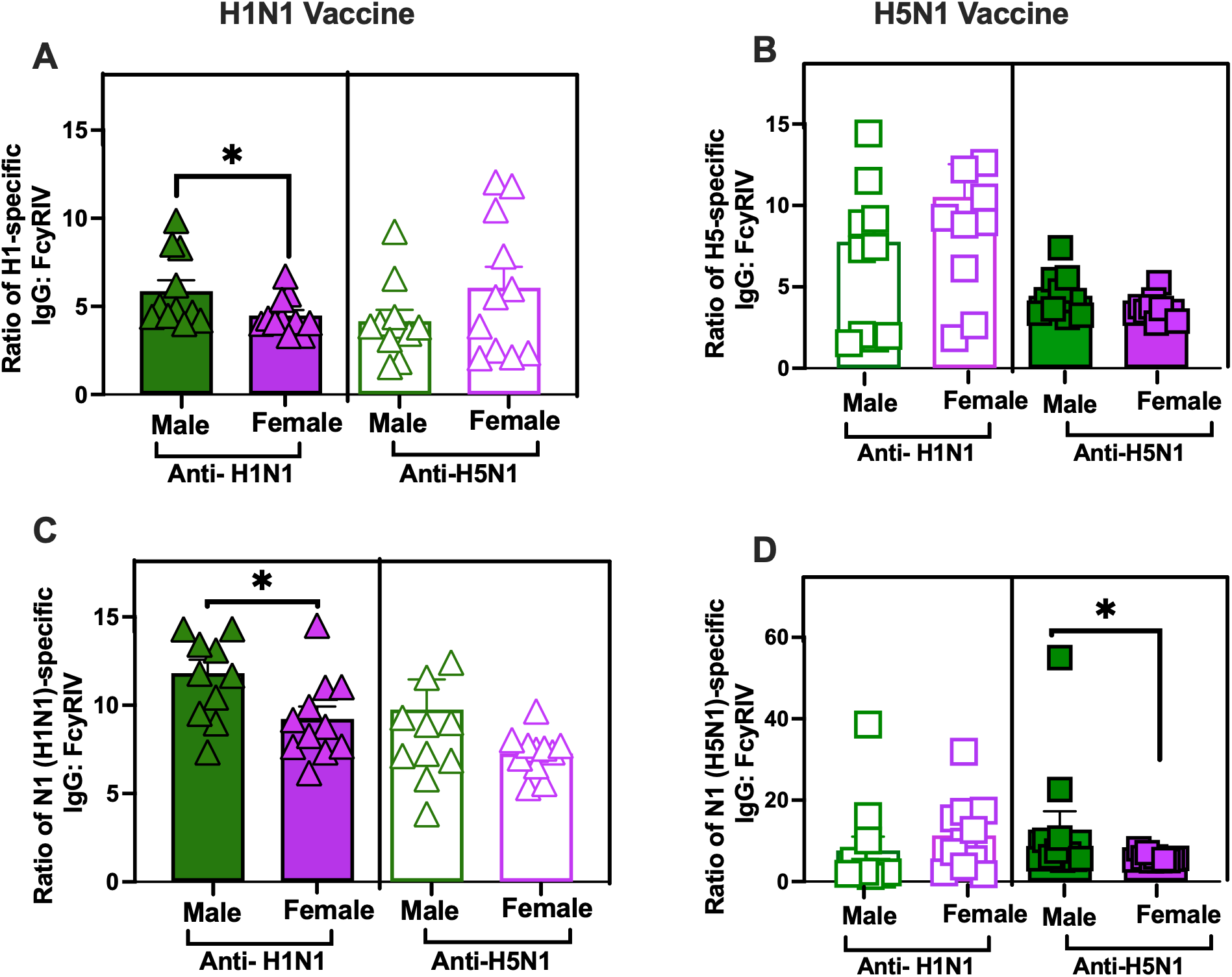
Antigen-specific IgG:FcγRIV ratios following H1N1 and H5N1 vaccination in C57BL/6 mice. Adult 7–8-week-old male (dark green) and female (dark purple) C57BL/6 mice were vaccinated twice, with a 3-week interval, using BPL-inactivated whole 2009 H1N1 or H5N1 (LAIV backbone) viruses in a prime/boost regimen. Plasma samples were collected 35 days post-vaccination (dpv) and antigen-specific IgG and FcγRIV binding were measured by multiplex systems serology. To account for differences in antigen-specific antibody quantity between the sexes, FcγRIV binding was normalized to antigen-specific IgG for each animal, with data shown as the ratio of antigen-specific IgG to FcγRIV binding. Ratios are shown for HA-specific responses in H1N1-vaccinated mice against homologous H1 and heterologous H5 antigens (A), HA-specific responses in H5N1-vaccinated mice against homologous H5 and heterologous H1 antigens (B), NA-specific responses in H1N1-vaccinated mice against homologous and heterologous N1 antigens (C), and NA-specific responses in H5N1-vaccinated mice against homologous and heterologous N1 antigens (D). Data are presented as mean ± SEM (n = 10– 11/group). Asterisks (*) indicate significant sex differences determined by Mann-Whitney tests, using GraphPad Prism 10.1.0, p<0.05.

**Fig. S3.**
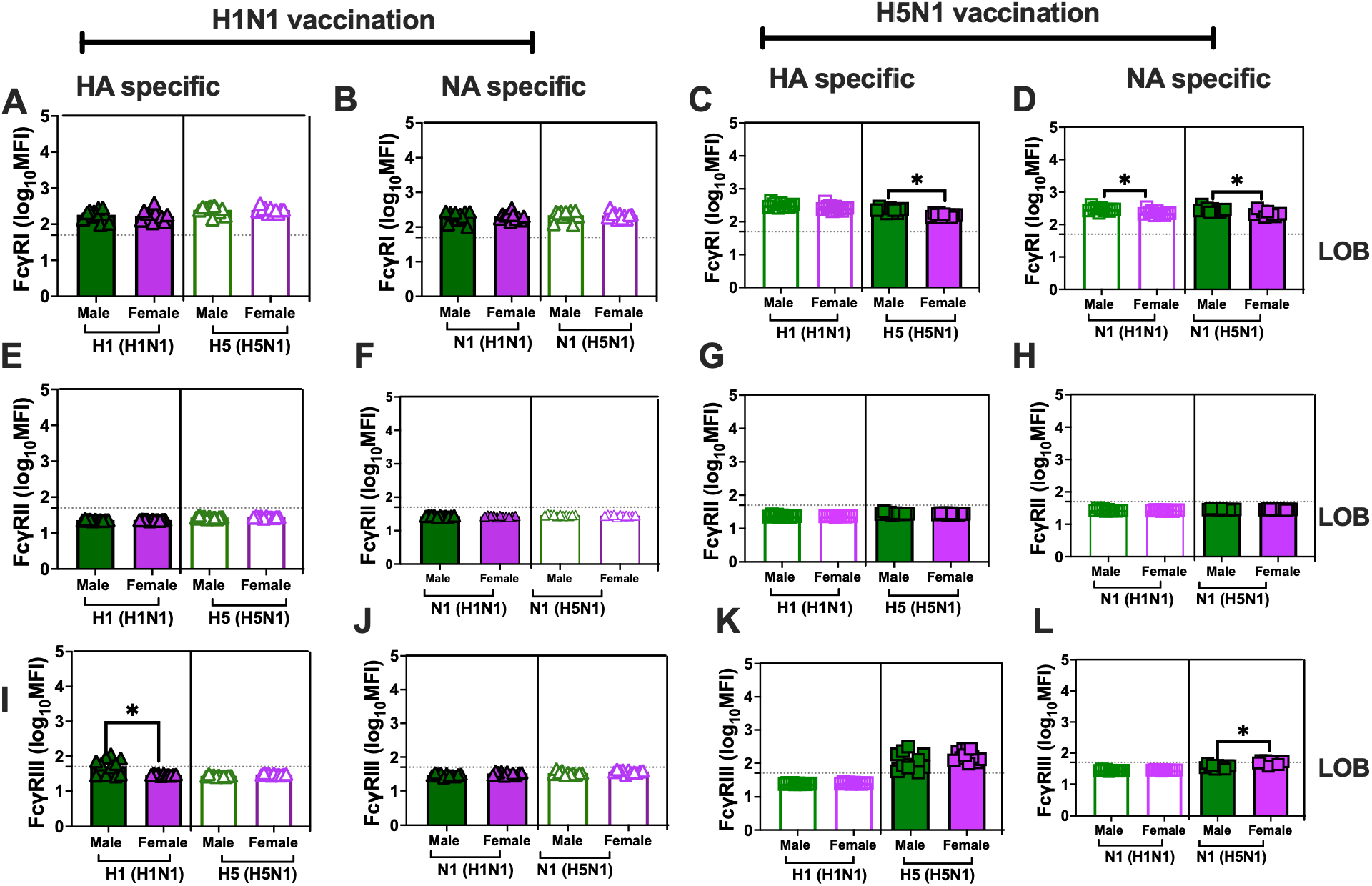
Fcγ receptor binding to selected HA and NA antigens following H1N1 and H5N1 vaccination in C57BL/6 mice. Adult 7–8-week-old male (dark green) and female (dark purple) C57BL/6 mice were vaccinated twice, with a 3-week interval, using BPL-inactivated whole 2009 H1N1 or H5N1 (LAIV backbone) viruses in a prime/boost regimen. Plasma samples were collected 35 days post-vaccination to determine antibody binding to FcγRI (A–D), FcγRII (E–H), and FcγRIII (I–L), measured by multiplex systems serology and shown as log10 mean fluorescence intensity (MFI) values for selected antigens, HA (H1 or H5) and NA (N1 from either H1N1 or H5N1). Data are presented as mean ± SEM (n = 10–11/group). Asterisks (*) indicate significant sex differences determined by Mann-Whitney tests, using GraphPad Prism 10.1.0. LOB = limit of background.

**Fig. S4.**
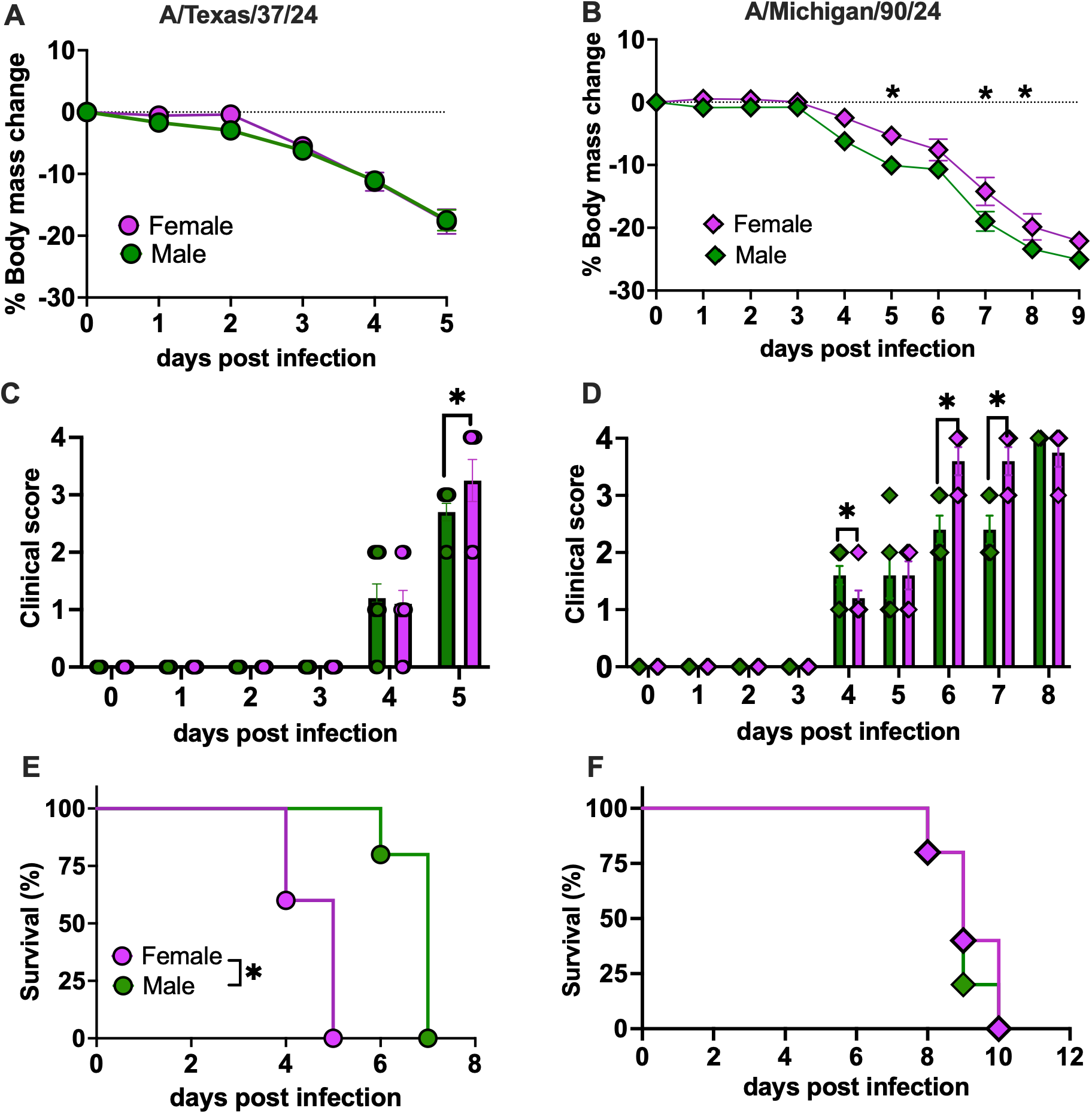
Morbidity and survival following A/Texas/37/2024 and A/Michigan/90/2024 H5N1 challenge in C57BL/6 mice. Adult 7–8-week-old male (dark green) and female (dark purple) C57BL/6 mice were intranasally inoculated with 10^2^ TCID_50_ of A/Texas/37/2024 (A, C, E) or A/Michigan/90/2024 (B, D, F). For the morbidity study, body mass changes (A, B), clinical scores (C, D), and survival (E, F) were recorded daily for up to 5 days post-infection for A/Texas/37/2024 and up to 9 days post-infection for A/Michigan/90/2024. Data are presented as mean ± SEM (n = 5–6/group). Asterisks (*) indicate significant sex differences (p < 0.05) as determined by two-way ANOVA using GraphPad Prism 10.1.0.

**Fig. S5.**
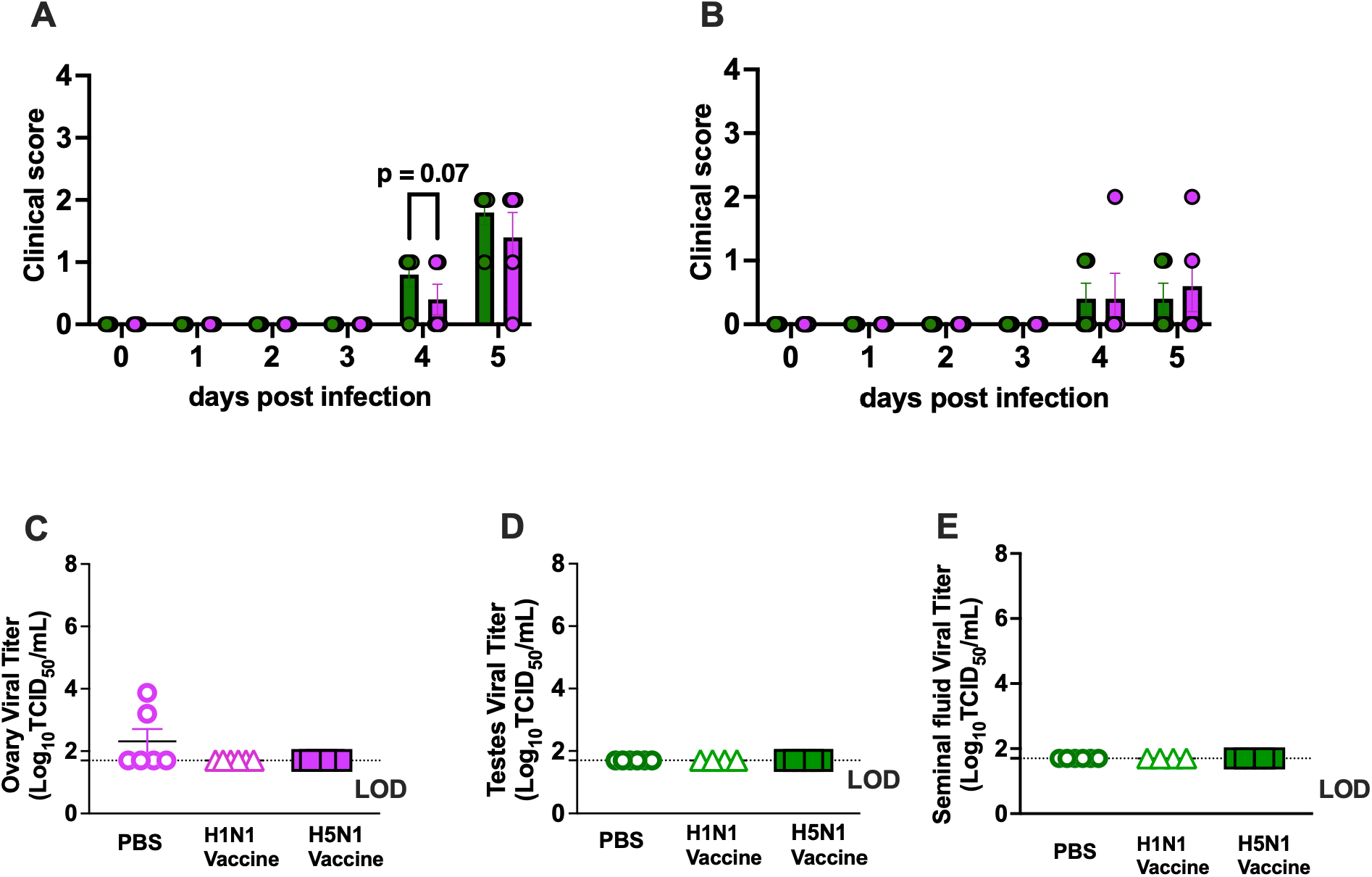
Clinical scores and extrapulmonary virus titers following A/Texas/37/2024 H5N1 challenge in vaccinated C57BL/6 mice. Adult 7–8-week-old male (dark green) and female (dark purple) C57BL/6 mice were vaccinated twice, with a 3-week interval, using BPL-inactivated whole 2009 H1N1 or H5N1 (LAIV backbone) viruses in a prime/boost regimen, and challenged intranasally with 10^2^ TCID_50_ of A/Texas/37/2024 at 48 days post-vaccination. Clinical scores were recorded daily for 5 days post-challenge in H1N1-vaccinated (A) and H5N1-vaccinated (B) mice (n = 5/sex). Infectious virus was quantified in ovary (C), testes (D), and seminal fluid (E) at 3 days post-challenge (n = 5/group). Asterisks (*) indicate significant sex differences (p < 0.05) as determined by two-way ANOVA using GraphPad Prism 10.1.0.

